# Role of Alanine Transaminase in Retinal Metabolic Homeostasis: Potential therapeutic target in retinal diseases

**DOI:** 10.64898/2026.04.19.719493

**Authors:** Qin Chen, Ting Zhang, Jingwen Zeng, Michelle Yam, Sora Lee, Fanfan Zhou, Meidong Zhu, Ming Zhang, Fang Lu, Jianhai Du, Mark Gillies, Ling Zhu

## Abstract

**Purpose:** Alanine transaminases (ALT), encoded by the GPT gene, catalyzes the reversible conversion of pyruvate and glutamate to alanine and alpha-ketoglutarate, thereby correlating carbohydrate and amino acid metabolism. However, its role in the human neural retina remains unclear. This study aimed to explore the expression, localization, and metabolic function of ALT in the human neural retina and its potential involvement in retinal diseases.

**Methods:** ALT1 and ALT2 expression and localization were examined in the retinas of healthy and diabetic retinopathy (DR) donors via immunoblotting and immunofluorescence. ALT function was assessed in *ex vivo* human retinal explants using pharmacological inhibition with beta-chloro-L-alanine (BCLA), followed by the analyses of enzyme activity, tissue injury, and transcriptomic responses. Stable-isotope tracing with ^13^C-and ^15^N-labelled substrates combined with GC-MS was used to define ALT-dependent carbon and nitrogen fluxes in macular and peripheral retinas. Redox level (NADPH/NADP^+^) was also evaluated under tert-butyl hydroperoxide-induced oxidative stress.

**Results:** ALT1 and ALT2 were both expressed in the human neural retina, with prominent localization in Müller glia and photoreceptor inner segments. ALT1 displayed a diffuse cytoplasmic distribution, whereas ALT2 demonstrated a punctate pattern consistent with mitochondrial localization. In DR retinas, ALT1 expression was spatially disorganized and heterogeneous, while ALT2 remained comparatively preserved. Inhibition of ALT with BCLA markedly reduced ALT activity without causing overt cytotoxicity or major transcriptional changes. Isotope tracing demonstrated that retinal ALT predominantly channels pyruvate-derived carbon into alanine, whereas alanine was minimally contributed to pyruvate production under basal conditions. ALT inhibition suppressed alanine synthesis and release, redirected nitrogen flux towards glutamate, glutamine, and aspartate, and uncovered distinct metabolic adaptations in macular but not peripheral retinas. Under oxidative stress, ALT inhibition induced the decrease of NADP^+^/NADPH ratio and LDH release, indicating improved redox balance and reduced tissue injury.

**Conclusions:** ALT is previously unrecognized as a regulator of carbon and nitrogen partitioner in the human neural retina, contributing to redox homeostasis under stress. The altered distribution of ALT1 in DR retina and the protective metabolic effects of ALT inhibition suggest ALT as a potential contributor to retinal metabolic vulnerability and a candidate therapeutic target in retinal diseases.

## Introduction

Alanine transaminase (ALT), encoded by the GPT gene, is an enzyme that reversibly converts pyruvate and glutamate into alanine and α-ketoglutarate (α-KG). It has two isoenzymes, ALT1 and ALT2, encoded by *Gpt1* and *Gpt2* ^1^. ALT1 is cytosolic and highly expressed in the liver, adipose tissues, skeletal muscles, and intestine, whereas ALT2 is a mitochondrial matrix enzyme enriched in metabolically active tissues including the skeletal muscles, heart, brain, and liver ^2,3^. Despite extensive characterization of their systemic roles, the distribution and function of ALT in the human retina remain poorly defined.

ALT correlates carbohydrate and amino acid metabolism, supporting carbon–nitrogen exchange, anaplerosis, and redox maintenance across multiple organs ^4^. Circulating ALT is widely used as a biomarker of hepatic injury ^5^. Furthermore, the relationship between ALT and diabetes mellitus (DM), a prevalent metabolic disease that seriously threatens global health, has been reported ^6,7^. Elevated ALT has been shown to be a predictor of type 2 DM ^8^. The expression of ALT2 in hepatocytes was also increased in diabetic conditions ^9^. In some diabetes-related complications, the S-shaped increase of ALT level was closely related to ketoacidosis-induced cell death ^10^. A U-shaped nonlinear association between ALT and diabetic kidney disease has also been demonstrated ^11^. These findings suggested ALT as a metabolic integrator and DM-associated enzyme, but its role in retinal metabolism has not been extensively explored.

Diabetic retinopathy (DR) affects over 20% of individuals with diabetes and remains a major cause of vision loss in working-age adults ^12^. DR arises from the convergence of vascular injury, inflammation, neuronal dysfunction, and metabolic stress. Dysregulated glucose metabolism is a principal driver of retinal vulnerability ^13^, and growing evidence implicates amino acid pathways including glutamate, proline, and branched-chain amino acid metabolism, in the disease progression of DR ^14,15^. Recent metabolomic analyses of vitreous and aqueous humour has identified pyruvate, alanine and glutamate as the key metabolites in DR ^13,16^, although plasma-based studies show incomplete concordance with ocular sample findings ^17^.

The relationship between DR severity and systemic ALT level has not been elucidated. It has been reported that ALT level was statistically different when comparing non-diabetic and DR groups, but no significant difference was found within DR groups at different disease stages ^18^. The results from the same study cohort via different methods show inconsistent correlations ^19^. Thus, it is important to elucidate ALT expression and function in the human retina.

Our study is the first comprehensive analysis of ALT1 and ALT2 expression, localization, and metabolic activity in healthy and DR retinas. Taking advantage of β-chloro-L-alanine (BCLA, a selective ALT inhibitor) ^20^ and ^13^C/^15^N isotope tracing, we explored the metabolic consequences of ALT inhibition on carbon and nitrogen fluxes, spatial metabolic organization, and oxidative stress responses in the unique *ex vivo* human retinal explant model. We observed distinct isoform-specific expression patterns of ALT. We also found that ALT actively shapes retinal metabolic networks. Such finding indicated that ALT is a previously unrecognized regulator of retinal redox balance and a potential therapeutic target in retinal metabolic diseases, particularly DR.

## Materials and Methods

### Ethics Approval and Human Donor Tissue

Human retinal tissues were obtained from the Lions New South Wales Eye Bank. All procedures adhered to the Declaration of Helsinki and were approved by the Human Research Ethics Committee at the University of Sydney (HREC#16/282) and the University of Sydney Institutional Biosafety Committee (21E011). Informed consent for research use was obtained for all donor tissues. Post-mortem intervals did not exceed 36 hours (Donor information in Supplementary Table 1).

### Frozen sections

Human retinas were fixed in 4% paraformaldehyde for 1 hour at room temperature, rinsed in PBS, and embedded in Optimal Cutting Temperature compound (ProSciTech, IA012). Samples were snap-frozen in liquid nitrogen and sectioned at 16 µm using a cryostat (Cryostar NX50, Epredia). Sections were stored at −30 °C until use.

### Immunofluorescence Staining

Frozen sections were defrosted to room temperature and blocked in PBS containing 10% normal donkey serum for 1 hour. Primary antibodies diluted in PBS with 1% donkey serum and 1% Triton X-100 were applied for 48 hours at 4 °C. After washing, sections were incubated with Alexa Fluor 488- or 594-conjugated secondary antibodies for 2 hours at room temperature. Nuclei were counterstained with Hoechst 33342. Slides were mounted with Vectashield antifade medium and imaged on a Zeiss LSM 700 confocal microscope. Image contrast and brightness were adjusted uniformly in ZEN software. Primary antibodies: ALT1 (Abcam, ab202083; 1:200), ALT2 (Abcam, ab101876; 1:200), CRALBP (Abcam, ab15051; 1:500).

### Western Blot

Retinal proteins were extracted in RIPA buffer (Sigma-Aldrich, R0278) containing protease/phosphatase inhibitors (Cell Signaling Technology, 5872S). Lysates were mixed with NuPAGE loading dye and reducing buffer (Life Technologies) and denatured at 70 °C for 10 minutes. Proteins were resolved on 4–12% Tris-Bis NuPAGE gels (Life Technologies) by electrophoresis at 180 V for 70 minutes at 4 °C. Transfer to PVDF membranes (Millipore) was performed at 100 V for 1.5 hours at 4 °C. Membranes were blocked in 5% BSA in TBST (0.1% Tween-20) for 1 hour and probed with primary antibodies overnight. After incubation with HRP-conjugated secondary antibodies, signals were detected using ECL reagents and quantified with Gene Tools.

Primary antibodies: ALT1 (Abcam, ab202083; 1:1000), ALT2 (Santa Cruz, sc-398383; 1:1000) and GAPDH (Cell Signaling Technology, 2118; 1:1000)

### Human retina explant dissection and culture

Donor eyes were stored in CO₂-independent medium at 4 °C. After removal of anterior structures, fundus imaging was performed and the neural retina was gently isolated. Vitreous was removed by blunt dissection. Five-millimeter explants were trephined from the macula and superior mid-peripheral retina and placed photoreceptor-side down on transwell inserts in the modified neuronal culture medium (The composition of the culture medium is based on Neurobasal™-A Medium, see Supplementary Table 2). Retinal explants were cultured at 37°C in 5% CO2 incubator.

### ALT Activity Assay

Five-millimeter human retinal explants were homogenized in ALT assay buffer, centrifuged at 12,000 g for 5 minutes at 4 °C, and supernatants were transferred to assay plates. ALT activity was quantified using a fluorometric ALT assay kit (Abcam, ab105134) with readings acquired every 3 minutes for at least 50 minutes (Ex/Em 535/587 nm).

### Lactate dehydrogenase (LDH) Cytotoxicity Assay

Conditioned culture medium collected after 24-hour explant incubation was processed using the Pierce LDH Cytotoxicity Kit (ThermoFisher, 88954). After 30 minutes at room temperature, absorbance at 492 nm was measured using a FLUOstar Omega plate reader.

### RNA sequencing

Retinal explants treated with 8 µM BCLA for 4 hours were snap-frozen at −80 °C. RNA was extracted using the Single-Cell RNA Purification Kit (Norgen, 51800) with on-column DNase digestion (QIAGEN, 79254). RNA concentration and integrity were confirmed using a Qubit RNA HS assay (Invitrogen, Q32852). Samples were sent to Novogene (Singapore) for bulk RNA-seq with 150 bp paired-end sequencing. Library quality was assessed with an Agilent Bioanalyzer 2100.

### Isotope labelling and metabolites extraction

We collected the macular and peripheral retinal explants from 5 pairs of post-mortem eyes for each labelling group. Retinal explants were incubated in medium with ^13^C-glucose (5mM, CLM-1396-0.5, Cambridge Isotope Lab), ^15^N-glutamine (2mM, 487809-500MG, Sigma-Aldrich), ^13^C-pyruvate (0.02mM, 490733-250MG, Sigma-Aldrich), or ^13^C-alanine (0.23mM, CLM-116-0.5, Cambridge Isotope Lab) for 4 hours. The basal medium used for the isotope tracing experiments was our modified neuronal culture medium (see Table 2). For each labelling experiment, the isotopically labelled compound was added by replacing the corresponding unlabelled component in the medium at an equimolar concentration (see Supplementary Table 3). Left-eye explants received 8 µM BCLA; right-eye explants served as controls. Metabolites were extracted from neuroretina using methanol:chloroform:water (700:200:50) and medium metabolites were collected in 80% methanol. Samples were dried and analyzed by gas chromatography mass spectrometry (GC-MS).

### NADP^+^/NADPH assay

Four-millimeter retinal explants punches were homogenized in 1% DTAB buffer and processed using the NADP^+^/NADPH-Glo Assay kit (Promega, G9081). Luminescence was recorded and ratios calculated as indicators of oxidative stress.

### Statistical Analysis

Data are presented as mean ± standard error of the mean. Paired two-tailed t-tests were used for two-group comparisons, and one-way ANOVA was applied for analyses involving more than two groups. Significance was set at p < 0.05. GraphPad Prism (version 10.0, GraphPad Software, CA) was used for all statistical analysis and data visualization.

## Results

### ALT expression and localization in normal and DR retinas

We first examined the expression of ALT1 and ALT2 protein in human post-mortem neural retinas. Both isoforms were consistently present in the retinal samples collected from multiple donors (Fig. S1). In comparison, the expression of ALT1 and ALT2 was only detected in the mouse (C57BL/6) RPE samples but not in the neural retina (Fig. S2).

Having confirmed their presence, we next sought to define the spatial distribution and cellular localization of ALT1 and ALT2 in human retina using immunofluorescence (IF) staining. Given the close association between ALT and DM, these analyses were performed in retinas from both non-diabetic donors and individuals with DR.

In the normal retinas (Fig. 1A-F), ALT1 exhibited cytoplasmic immunofluorescent staining throughout Müller glial cells, extending from their endfeet at the inner limiting membrane to their outer processes. This cytoplasmic localization was further confirmed by the colocalization with Cellular Retinaldehyde Binding Protein (CRALBP, a known cytoplasmic marker of Müller glia) (Fig. 1C). The most significant colocalization was observed in the main cell bodies within the inner nuclear layer (INL) and the radial processes adjacent to the ganglion cell layer (GCL). ALT1 was also detectable within the inner segments of photoreceptors. In contrast, ALT2 showed a distinct pattern (Fig. 1D, F). Although ALT2 was present throughout Müller glia, its staining appeared punctate and granular, consistent with its known mitochondrial localization. A prominent ALT2 signal was also observed in photoreceptor inner segments.

**Figure 1:**
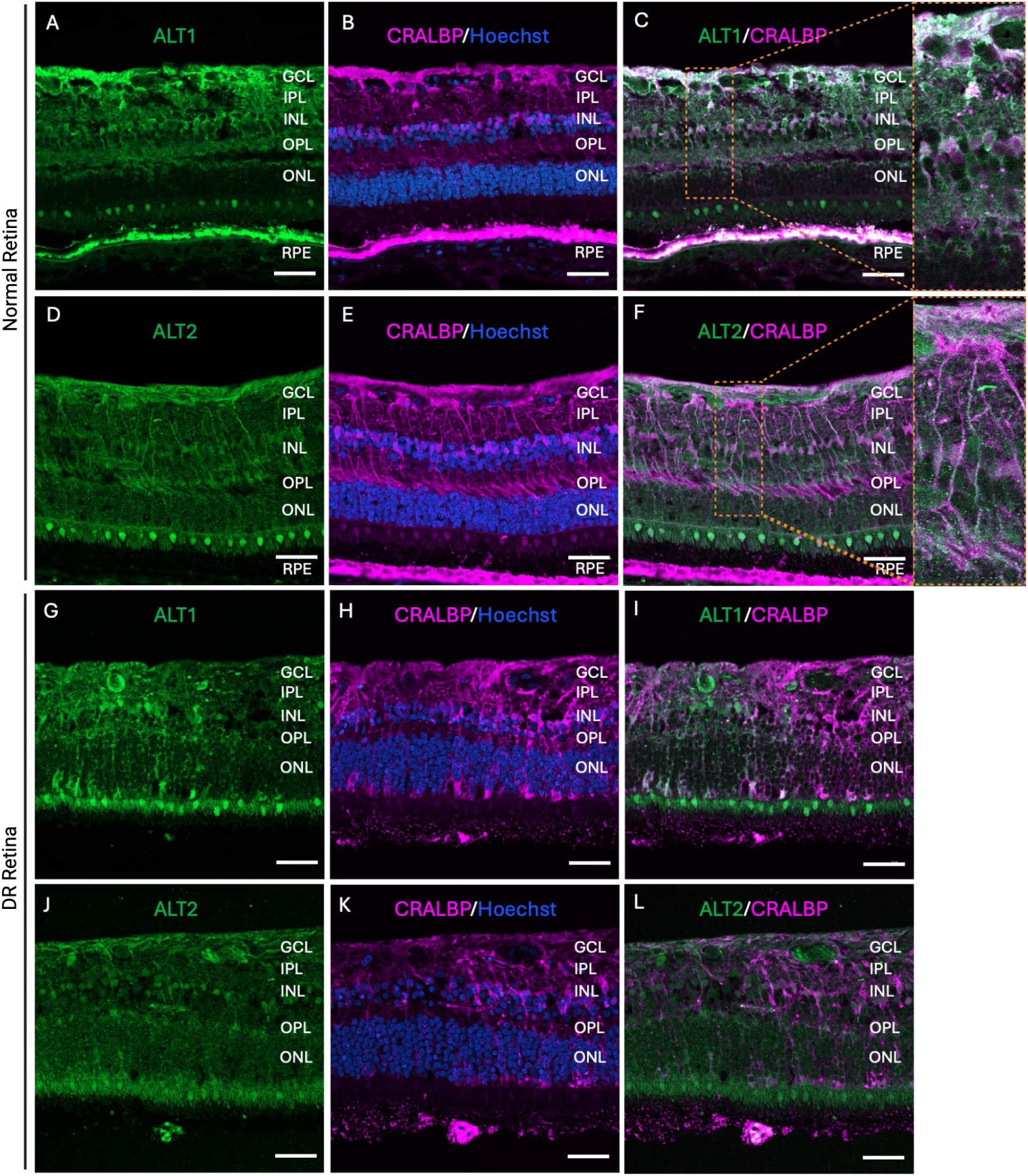
ALT1 and ALT2 show distinct localization patterns in human retinas and differential alterations in diabetic retinopathy. Confocal immunofluorescence images of human retinal sections showing the localization of ALT1 and ALT2 in non-diabetic and DR retinas. ALT1 or ALT2 is shown in green, CRALBP in magenta marks Müller glia, and Hoechst in blue labels nuclei. A-F: In the normal peripheral retinas, both ALT1 (A-C) and ALT2 (D-F) were detected throughout Müller glial cells, extending from the endfeet at the inner limiting membrane to the outer radial processes. Both isoforms were present in photoreceptor inner segments. G-L: In the DR retinas, ALT1 staining are spatially heterogeneous, with some regions with increased ALT1 signal corresponding to areas of reduced CRALBP intensity (G-I), indicating disruption of Müller glial ALT1 distribution. In contrast, ALT2 staining remained comparatively consistent in Müller glia and photoreceptor inner segments (J-L). Scale bar = 50 μm. Key: GCL, ganglion cell layer; IPL, inner plexiform layer; INL, inner nuclear layer; OPL, outer plexiform layer; ONL, outer nuclear layer; RPE, retinal pigment epithelium.

In the retinas of DR patients (Fig. 1G-L), ALT1 expression showed a marked spatial disruption. Müller glial staining became patchy and heterogeneous around the peripheral retinas, with several regions of intensified ALT1 staining alternated with areas of attenuation. Notably, this dysregulation demonstrated an inverse relationship with CRALBP expression in peripheral retinas; the regions with reduced CRALBP exhibited elevated ALT1. However, ALT1 expression in photoreceptor inner segments remained relatively preserved. Conversely, the pattern of ALT2 staining in Müller glial cells remained consistent in both the normal and DR retinas (Fig. 1J-L).

### Metabolic Role of ALT in Human Neuronal Retinas

Upon validating the expression of ALT1 and ALT2 in the neuronal retinas, we then explored the role of ALT in human retinal metabolism. We adopted a pharmacological inhibition strategy to functionally probe ALT-dependent metabolic pathways in the intact human neuronal retinal tissues. It is known that β-chloro-L-alanine (BCLA) is an ALT inhibitor. Upon the treatment of BCLA, the enzymatic activity, tissue viability, and transcriptome profile were assessed in human retinal explants.

Dose-response experiments were performed using retinal explants from three donors that were exposed to different BCLA concentrations (i.e. 0, 0.125, 1, 8, and 64 μM) for 4 hours. BCLA treatment resulted in a dose-dependent suppression of ALT enzymatic activity. Notably, BCLA treatment at 8 µM reduced ALT activity to approximately 10% of that of the control (p < 0.05) (Fig. S3A). Despite the substantial reduction of ALT activity, no significant increase in LDH release was observed across all concentrations, indicating that BCLA did not induce significant cytotoxicity under the current experimental conditions (Fig. S3B). Together, BCLA treatment at 8 μM was selected for the subsequent metabolic analyses as it was effective to inhibit ALT activity while preserving retinal explant viability.

To evaluate whether ALT inhibition induced broader transcriptional responses, we performed bulk RNA sequencing on macular and peripheral explants obtained from five donors with or without the treatment of BCLA. More than 30,000 transcripts were detected across all samples. Differential expression analysis identified 146 genes in macular explants (Fig. S4A, p < 0.05) and 195 genes in peripheral explants (Fig. S4B, p < 0.05). However, none remained significant after Benjamini-Hochberg correction (adjusted p > 0.1), and no apoptotic pathways were enriched using Interpretative Phenomenological Analysis.

To evaluate the impact of BCLA on the metabolic balance of alanine and pyruvate, we incubated retinal explants with either ¹³C-labelled pyruvate or ¹³C-labelled alanine for 4 hours to trace the changes of the metabolic fluxes upon ALT inhibition. The incorporation of ¹³C into M1-labelled alanine or pyruvate was then quantified. Using ¹³C-pyruvate as a tracer, 3.01% ± 0.64% of alanine in the macular and 4.15% ± 0.90% in the periphery retinas were labelled within 4 hours. Following the treatment of BCLA, ^13^C-labelled alanine was significantly reduced to 47.4% of the control in the macular (3.01% ± 0.64% *vs* 1.43% ± 0.21%, p = 0.037) and to 46.0% of the control in the periphery retinas (4.15% ± 0.90% *vs* 1.91% ± 0.38%, p = 0.016). Other labelled metabolites related to downstream pyruvate metabolic pathways did not show significant differences (Fig. 2A-D). In contrast, when ¹³C-alanine was used as the tracer, a minimal labelling of pyruvate was detected (0.39% ± 0.05% in the macular and 0.33% ± 0.04% in the periphery retinas). The labelled pyruvate did not show significant change with or without BCLA treatment (Fig. 2E and F).

**Figure 2:**
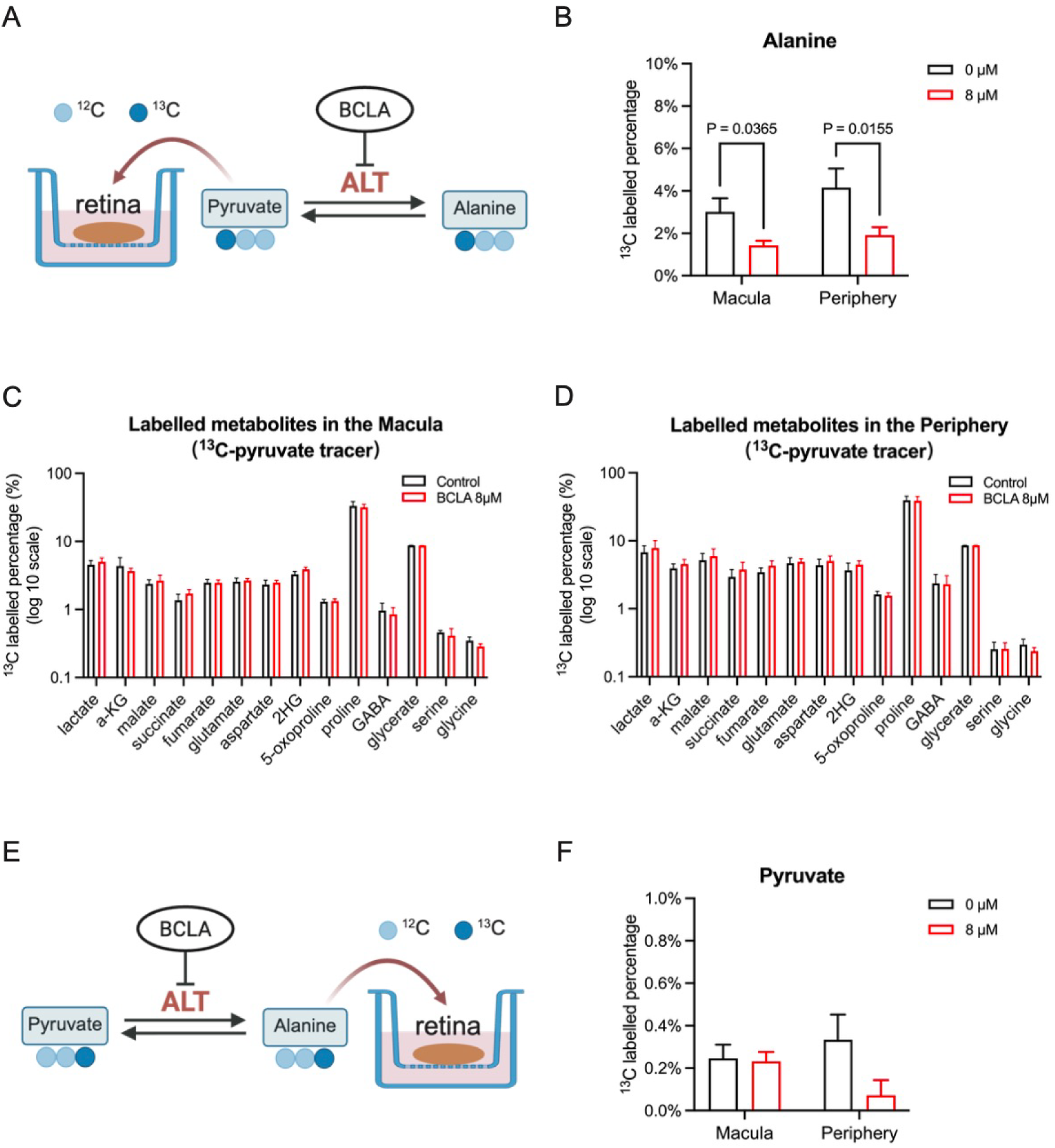
ALT inhibition suppresses pyruvate-to-alanine flux but has minimal influence on alanine-to-pyruvate flux in human retinal explants. Human macular and peripheral retinal explants from five donors were incubated with ^13^C-labelled pyruvate or ^13^C-labelled alanine for 4 h, in the presence or absence of BCLA. A & E: Schematic overview of the isotope-tracing strategy used to assess bidirectional ALT-dependent flux between pyruvate and alanine. B: When ^13^C-pyruvate was used as the tracer, BCLA significantly reduced the formation of ^13^C-labelled alanine in both macular and peripheral explants. C-D: The level of other pyruvate-derived metabolites was not significantly altered with or without BCLA treatment in either retinal region. F: When ^13^C-alanine was used as the tracer, only minimal labelling of pyruvate was detected with a minimal impact of BCLA treatment. Data are presented as mean ± SE.

### The effect of ALT inhibition on carbon and nitrogen metabolism

As pharmacological inhibition of ALT altered the metabolic flow from pyruvate to alanine, we then explored how ALT inhibition regulates carbon- and nitrogen-involved metabolic pathways in the neuronal retinas.

We incubated retinal explants with ¹³C-labelled glucose for 4 hours to trace the carbon flux in glucose metabolism. We found that BCLA treatment resulted in an approximately 70% reduction in ¹³C-labelled alanine in both the macular (55.5% ± 5.3% vs 14.6% ± 3.8%, p < 0.001) and peripheral retinas (56.0% ± 7.3% vs 17.9% ± 4.5%, p < 0.001) (Fig. 3C). However, ¹³C-labelled pyruvate level remained unchanged with or without BCLA treatment in both the macular and periphery retinas (Fig. 3B). Downstream pyruvate-derived metabolites including lactate, α-KG, 2HG, malate, succinate, and fumarate, also showed no significant changes (Fig. 3D-I).

**Figure 3:**
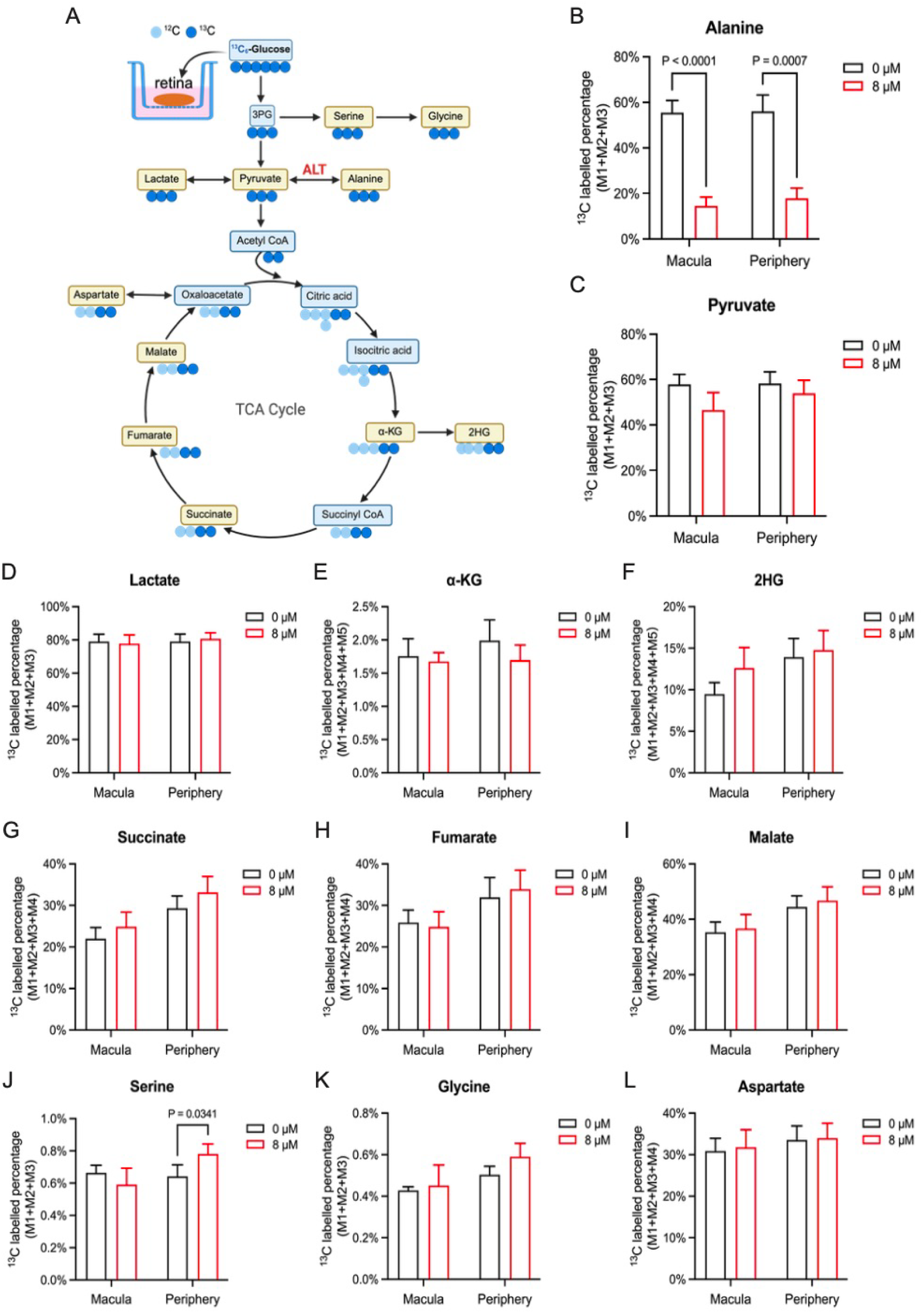
Altered glucose-derived carbon flux in human macular and peripheral retinal explants upon BCLA treatment. Human macular and peripheral retinal explants were incubated with ^13^C-glucose for 4 h in the presence or absence of 8 μM BCLA to evaluate the effect of ALT inhibition on glucose-derived carbon metabolism. A: Schematic diagram of ^13^C-glucose-derived carbon flow. Yellow boxes indicate metabolites measured in this study. B-L: Relative abundance of ^13^C-labelled metabolites in macular and peripheral explants with or without BCLA treatment. ALT inhibition resulted in a marked reduction in ^13^C-labelled alanine in both macular and peripheral retinas, while ^13^C-labelled pyruvate remained unchanged. Most downstream pyruvate-derived metabolites, including lactate, α-KG, malate, succinate, and fumarate, were not significantly altered. Data are presented as mean ± SE.

Given the coupling of alanine metabolism and other *de novo* amino acid synthesis^21^, we also examined the impact of ALT inhibition on serine-glycine metabolism, which is very important in the one-carbon metabolism in the neural retina^22^. With an exogenous supply of serine and glycine, ALT inhibition significantly increased the level of labelled serine in the periphery (0.64% ± 0.06% *vs* 0.78% ± 0.07%, p = 0.034), whereas glycine level was not significantly altered in both macular and periphery retinas (Fig. 3J and K).

We also extracted the labelled metabolites in the culture to assess how the releases of metabolites upon ALT inhibition (Fig. S5). The results showed a reduction in ^13^C-alanine release from both macular (31.1% ± 4.4% *vs* 19.9% ± 3.6%, p < 0.001) and peripheral (28.2% ± 2.4% *vs* 17.3% ± 2.8%, p = 0.005) explants upon the exposure to BCLA (Fig. S5B). Such findings suggested that ALT activity contributes not only to intracellular alanine pools but also to alanine export by the retina. A slight increase was observed in the release of other metabolites from the macular region including fumarate (34.9% ± 2.4% *vs* 41.8% ± 3.8%, p = 0.065), malate (16.1% ± 3.1% *vs* 19.3% ± 2.8%, p = 0.072), serine (0.87 ± 0.08% *vs* 1.05 ± 0.11%, p = 0.087), and glycine (0.87 ± 0.05% *vs* 0.96 ± 0.04%, p = 0.009) (Figure S5G-J).

To trace the nitrogen flux during amino acid metabolism, we incubated retinal explants with ^15^N-labelled glutamine for 4 hours. Compared with the control group, BCLA treatment reduced the level of ^15^N-labelled alanine by approximately 76% in the macular (8.6% ± 1.1% *vs* 35.8% ± 4.4%, p < 0.001) and by approximately 70% in the peripheral retina (7.9% ± 1.5% *vs* 26.1% ± 6.7%, p = 0.027) (Fig. 4B). ^15^N-labelled glutamate and glutamine levels showed significant increases in both macular and peripheral regions (Fig. 4C and D). As to the nitrogen flux redirection from glutamate upon BCLA treatment, the level of labelled aspartate was significantly elevated by approximately 21% in the macular (54.0% ± 1.4% *vs* 44.8% ± 1.9%, p = 0.005) and by approximately 11% in the periphery (48.9% ± 5.2% *vs* 44.0% ± 5.6%, p = 0.036) (Fig. 4E). Conversely, ^15^N-labelled GABA was reduced to approximately 77% of the control in the macular (3.69% ± 0.37% *vs* 2.84% ± 0.09%, p = 0.042) and to approximately 85% of the control in the periphery (3.26% ± 0.20% *vs* 2.76% ± 0.06%, p = 0.041) following BCLA treatment (Fig. 4G). Other glutamate-derived metabolites including proline, ornithine, serine, and glycine, were not significantly altered (Fig. 4H-K).

**Figure 4:**
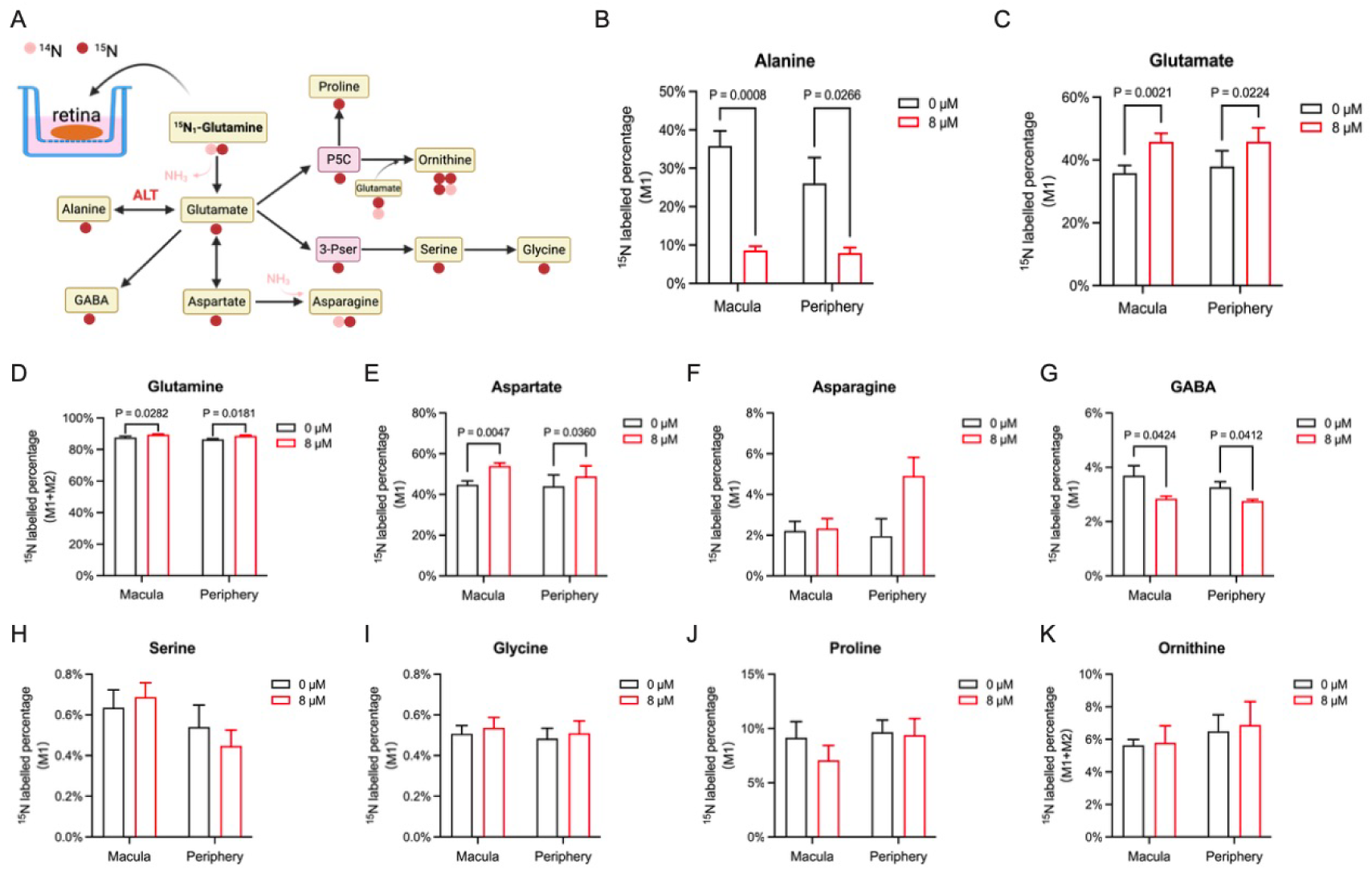
Changes of glutamine-derived nitrogen flux in retinal explants following BCLA treatment. Human macular and peripheral retinal explants were incubated with ^15^N-glutamine for 4 h with or without 8 μM BCLA treatment to assess the contribution of ALT to nitrogen redistribution. A: Schematic diagram of ^15^N-glutamine-derived nitrogen flow into alanine and related amino acid pools. Yellow boxes indicate metabolites measured in this study. B-J: Percentage of ^15^N-labelled metabolites in macular and peripheral explants with or without BCLA treatment. BCLA markedly reduced ^15^N incorporation into alanine in both retinal regions. In parallel, ^15^N-labelled glutamate and aspartate increased significantly, indicating a re-route of nitrogen flux towards alternative transamination pathways. ^15^N-labelled GABA was significantly reduced, whereas proline, ornithine, serine, and glycine were not significantly changed. Data are presented as mean ± SE.

### Regional metabolic heterogeneity following ALT inhibition

We also quantified the total abundance of metabolites in the culture collected from the explants of all 15 donors with or without 8 µM BCLA treatment using ^13^C tracing to explore the differential roles of ALT on the metabolism in the macular and periphery neural retinas. These analyses revealed a pronounced regional heterogeneity in metabolic responses to ALT inhibition.

In the macula, ALT inhibition led to ∼23% reduction in total alanine abundance (log2 fold change = −0.38, p = 0.001) and ∼21% reduction in pyruvate abundance (log2 fold change = −0.33, p < 0.001). In contrast, several metabolites reduced in the macula exhibited an opposite pattern in the peripheral retinas. Following ALT inhibition, the level of lactate, α-ketoglutarate, glutamine, 5-oxoproline, ornithine, malate, serine, glycerate, and 2-hydroxyglutarate was increased to approximately 110% ∼ 135% of the control in peripheral explants, while decreased in the macular region. Notably, GABA level was significantly increased by approximately 37% of the control in the peripheral retinas (log2 fold change = 0.46, p = 0.0114).

### ALT inhibition reduces the ratio of NADP^+^/NADPH

ALT correlates glucose- and amino acid-derived carbon and nitrogen fluxes, processes that are intrinsically coupled to cellular redox homeostasis. Given that oxidative stress is a key pathogenic driver of DR, ALT1 expression was increased in the retinas of DR. We then investigated whether ALT activity modulates redox status in the neuronal retinas.

We established an oxidative stress model of human retinal explants with the NADP^+^/NADPH ratio as an indicator. The retinal explants were exposed to increasing concentrations of tert-butyl hydroperoxide (TBHP) to induce oxidative stress, and oxidative burden was validated with the ratio of NADP^+^/NADPH. In the control explants, the ratio of NADP^+^/NADPH was 0.51 ± 0.11. Exposure to 800 μM TBHP, this was significantly increased to 1.32 ± 0.17 (p = 0.0065), indicating the induction of a substantial oxidative stress. In contrast, 400 μM TBHP did not significantly alter the ratio of NADP^+^/NADPH (0.45 ± 0.02, p = 0.569) (Fig. 5B).

**Figure 5:**
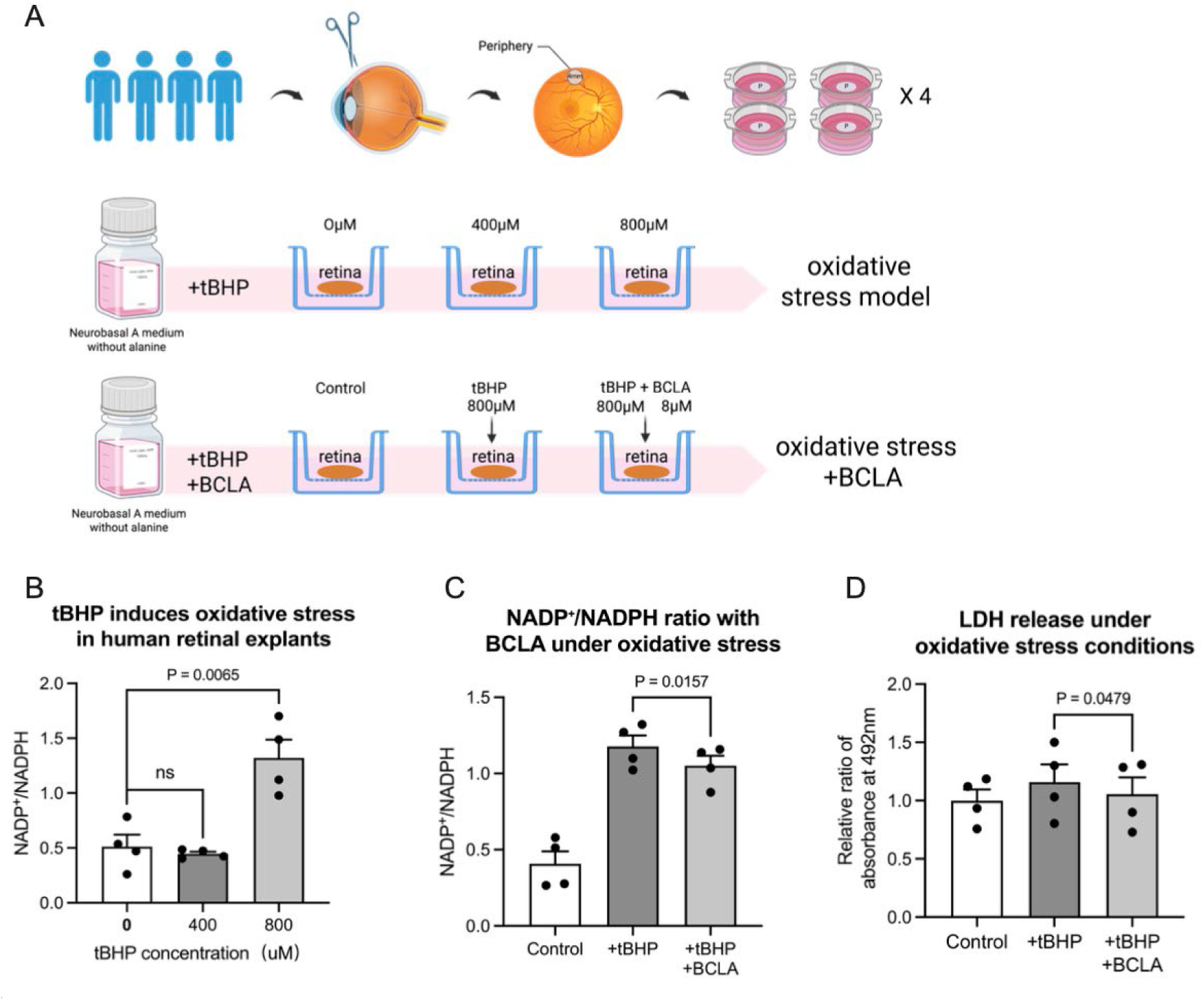
ALT inhibition reduces the ratio of NADP^+^/NADPH and tissue injury under oxidative stress in human retinal explants. The impact of ALT inhibition on retinal redox status was assessed in human retinal explants upon tert-butyl hydroperoxide (TBHP)-induced oxidative stress. A: Experimental design showing induction of oxidative stress with TBHP and the co-treatment of BCLA. B: Exposure to 800 μM TBHP for 4 h significantly increased the ratio of NADP^+^/NADPH, proving an induction of oxidative stress. C: Under oxidative stress, 8 μM BCLA treatment significantly reduced the ratio of NADP^+^/NADPH. D: LDH release, an indicator of tissue injury/cytotoxicity, was significantly reduced in the explants treated with TBHP plus BCLA compared with TBHP alone. Data are presented as mean ± SE.

Under TBHP-induced oxidative stress, an inhibition of ALT activity significantly altered redox balance. The treatment of 8 μM BCLA reduced the ratio of NADP^+^/NADPH from 1.22 ± 0.08 to 1.03 ± 0.05 (p = 0.0175), indicating a partial attenuation of oxidative stress (Fig. 5C). Consistently, LDH release, a marker of cytotoxicity, was also significantly reduced in the TBHP + BCLA group compared with TBHP alone (decreasing around 10% of the control, p = 0.0479), suggesting that BCLA treatment alleviates oxidative stress-induced cellular damage (Fig. 5D).

## Discussion

In this study, we provide the first comprehensive characterization of ALT isoforms in the human retina, revealing their involvement in retinal metabolic homeostasis and suggesting its association with metabolic dysregulation in DR. While ALT is widely recognized as a systemic metabolic regulator and biomarker for liver injury, its specific role in the retina was not well understood. Our findings showed that ALT is not merely present in the retina but is actively engaged in regulating carbon and nitrogen fluxes, with a distinct isoform localization disrupted in the DR retinas. Furthermore, an inhibition of ALT activity reprograms glucose and amino acid metabolism as well as attenuating oxidative stress, suggesting that ALT may represent an important metabolic regulator influencing retinal responses to metabolic stress.

Our immunofluorescence staining demonstrated that both ALT1 and ALT2 were prominently expressed throughout the Müller glial processes and cell bodies as well as in photoreceptor inner segments (Fig. 1). This localization aligns with the high glucose and glutamate metabolic demands of these cell types ^23–25^. Given the central role of Müller glia in neurotransmitter recycling, metabolic support and maintenance of neuronal homeostasis ^26,27^, the diffused cytoplasmic presence of ALT1 and the punctate mitochondrial localization of ALT2 suggest that ALT isoenzymes likely facilitate the shuttle of carbon and nitrogen between cellular compartments, similar to their roles in other high-energy tissues like the skeletal muscle and brain^1–3^. Meanwhile, the previously reported biochemical parameters suggested that ALT1 (Km, ala = 34 mM) and ALT2 (Km, ala = 2 mM) exhibit a preference towards alanine anabolism and catabolism, respectively ^2,28^. Consistent with their distinct kinetic properties and subcellular localization, ALT1 and ALT2 may perform complementary roles in the retina. Cytoplasmic ALT1 may preferentially support alanine synthesis and its integration into the alanine-glucose cycle for high glycolytic flux, whereas mitochondrial ALT2 with a higher affinity for alanine, may facilitate alanine catabolism, thereby contributing to mitochondrial nitrogen handling and redox balance.

One notable observation from the current study is that ALT1 but not ALT2, undergoes marked dysregulation in the DR retinas (Fig. 1). ALT1 exhibited a patchy and dysregulated expression pattern, inversely correlated with the Müller glia marker CRALBP in peripheral regions. In DM, Müller glial cell function is impaired, shifting from a homeostatic and neuroprotective state to a reactive and pro-inflammatory phenotype, contributing to retinal edema, vascular leakage, and neurodegeneration ^29,30^. Thus, the altered ALT1 pattern observed in this study may reflect metabolic disturbances associated with Müller glial dysfunction in DR. Moreover, the localization of photoreceptor-associated ALT1 remains unchanged, highlighting a cell type-specific susceptibility. Although this observation was derived from a single donor DR sample, it is plausible that ALT1 may serve as an indicator of metabolic alterations in Müller glia in the disease progression of DR.

Our isotope tracing experiments revealed that retinal ALT primarily catalyzed the forward transamination of pyruvate and glutamate to generate alanine and α-ketoglutarate (Fig.2). We observed a substantial incorporation of ^13^C-pyruvate into alanine, which was robustly suppressed by ALT inhibition. Conversely, ^13^C-alanine minimally contributed to pyruvate pools, indicating that ALT in human retinal tissues predominantly operates in the forward direction and minimally to alanine catabolism under physiological conditions. Studies in animals have shown that glucose was metabolized in the glial cells, which supply alanine to the neurons, and then ammonium returns to the glia ^31^. Therefore, based on the current findings, ALT in human Muller cells may play a crucial role in alanine-glucose metabolism in glial cells and the retinal photoreceptor network.

We found that BCLA significantly decreased alanine synthesis from pyruvate and reduced alanine release into the culture medium, confirming that alanine in the neuronal retinas is predominantly biosynthesized from glucose-derived pyruvate via the ALT pathway. The downstream metabolites of glucose, lactate and TCA intermediates, were slightly increased, suggesting that ALT inhibition might promote pyruvate flux rather than obstruction. Inhibiting alanine biosynthesis, one of carbon’s metabolic pathways, may boost flux through other carbon pathways (Fig. 3). The current findings are in line with the previous report that loss of ALT activity inhibited the production of alanine in cancer cells while enhancing mitochondrial metabolism ^32^.

Interestingly, ALT inhibition selectively enhanced serine labelling in the peripheral retinas without increasing the glycine level, possibly reflecting compensatory activation of the one-carbon cycle (Fig. 3). One-carbon metabolism is a central metabolic pathway critical for the biosynthesis of several amino acids, methyl group donors, and nucleotides ^33^, which serves to replenish NADPH and maintain redox equilibrium ^34^. Serine can be catalyzed by serine palmitoyl transferase (SPT) to form sphingolipids, which support neuronal structure and cell signaling. When the serine level is low, alanine can combine with SPT to generate deoxysphingolipids, which are toxic to photoreceptors. Low serine is likely the underlying cause of Macular Telangiectasia Type 2 (MacTel2) ^35^. *In vitro* experiments have also confirmed that reducing the serine/alanine ratio significantly increased the level of harmful deoxysphingolipids ^36^. Patients with MacTel2 have a much higher prevalence of Type 2 DM than the general population ^37^. Some metabolomic studies suggested that serine metabolism is significantly disrupted in DR ^38,39^. For example, vitreous metabolomic analysis reported significantly a lower L-serine level in the vitreous of non-proliferative DR patients than that of healthy or diabetic eyes without retinopathy ^39^. Thus, the simultaneous reduction in alanine and increase in serine upon ALT inhibition may be a potential therapeutic approach for DR or MacTel2 that to be further explored in the future.

The ^15^N-glutamine tracing experiments revealed that ALT inhibition disrupts nitrogen redistribution across key amino acids (Fig. 4). In the retina, alanine likely functions as a terminal nitrogen sink, as nitrogen incorporated into alanine is not readily redistributed to other amino acids. Previous studies have shown that alanine is poorly converted into other amino acids even when exogenously supplied ^21^. Consistently, inhibition of alanine synthesis led to a significant accumulation of glutamate and glutamine, accompanied by a diversion of nitrogen flux towards aspartate. This “shunting” effect suggests that when ALT is dysregulated, the retina rely on alternative transaminases, likely Aspartate Transaminase (AST), to manage amino group transfer. The concurrent increase in aspartate may also facilitate mitochondrial anaplerosis, buffering TCA cycle intermediates and preventing energetic collapse ^40^. Interestingly, we also observed a reduction in GABA synthesis flux following ALT inhibition. Since glutamate is the direct precursor for GABA, this reduction might indicate that ALT supports nitrogen exchange within glutamate-GABA cycling. Glutamate is the excitatory neurotransmitter, while GABA is the primary inhibitory neurotransmitter. The precise balance between glutamatergic excitation and GABAergic/glycinergic inhibition is essential for maintaining the homeostasis and dynamic range required for the accurate phototransduction and photoelectric signal conversion across varying light conditions ^25,41^. Thus, our findings suggested that ALT modulates not only metabolic but also neurotransmitter homeostasis in the retina.

Our data also indicated a regional metabolic heterogeneity between the macular and peripheral retinas. ALT inhibition in the macula led to a significant reduction in the total abundance of alanine and pyruvate, whereas no significant changes were observed in the peripheral retinas. In contrast, several TCA-related metabolites in the peripheral region showed an upward trend following ALT inhibition (Fig. S6), whereas similar metabolites in the macular region tended to decrease. Although these changes were not statistically significant, they may reflect region-specific metabolic adaptations to ALT blockade. The macula, characterized by a high density of cone photoreceptors, elevated oxygen consumption, and greater metabolic activity ^42^, likely requires more active coordination of glycolytic metabolism and nitrogen recycling. Consequently, when ALT activity is inhibited, the macula may undergo distinct metabolic redistribution compared with the peripheral retinas to sustain the metabolic demands. Such spatial variation in metabolic resilience has been documented for other key enzymes such as phosphoglycerate dehydrogenase and pyruvate kinase ^43,44^. Notably, the total GABA abundance level was significantly increased in the peripheral retinas in contrast to the reduced ^15^N-GABA flux observed in tracing experiments, suggesting that the periphery retinas may compensate for altered nitrogen flow by adjusting steady-state neurotransmitter pools. Our results elaborate on the role of ALT in this process, emphasizing the need to consider retinal topography when targeting metabolic enzymes therapeutically.

Finally, we found that ALT inhibition enhanced retinal redox resilience (Fig. 5). Under TBHP-induced oxidative stress, ALT inhibition reduced the ratio of NADP⁺/NADPH and decreased LDH release, indicating an attenuation of oxidative burden and cytotoxicity. NADPH plays a central role in maintaining glutathione in its reduced form, which is essential for antioxidant defense in retinal tissues. The increased NADPH availability observed following ALT inhibition may arise from several metabolic adaptations including enhanced flux through one-carbon metabolism, altered malate-associated pathways or increased activity of the pentose phosphate pathway ^45,46^. Although these mechanisms remain to be further confirmed, they provide possible routes through which ALT inhibition may promote redox homeostasis. In addition, through inhibiting ALT, the retina may spare glutamate, a critical precursor for GSH synthesis, thereby enhancing its antioxidant capacity^47^.

Oxidative stress is widely recognized as a major driver of DR, largely due to hyperglycemia-induced overproduction of mitochondrial reactive oxygen species ^48^. Notably, we observed altered ALT1 expression in the DR retinas. Together with the metabolic and redox effects observed following ALT inhibition, these findings indicate that ALT-mediated metabolic regulation may influence retinal responses to diabetic metabolic stress. Thus, modulating ALT activity may be a complementary approach to existing antioxidant therapies.

Our study has some limitations. Although retinal explants preserve key tissue architecture and cell-cell interactions, it is acknowledged that the isotope tracing experiments were performed under *ex vivo* culture conditions. The nutrient composition of the culture medium including the availability and relative concentrations of metabolic substrates and supplements, cannot precisely mimic physiological conditions in the human neuroretina. At present, detailed quantitative information describing the *in vivo* nutritional supply and metabolic microenvironment of the human neuroretina remains limited, making it difficult to replicate native conditions in an experimental setting. Nevertheless, despite these constraints, the consistent metabolic remodeling induced by BCLA-mediated ALT inhibition supports that ALT plays a functional role in regulating metabolic homeostasis in human neuroretina. In addition, while BCLA selectively inhibits ALT, off-target effects on other transaminases cannot be excluded ^49^. Future studies employing genetic knockdown or isoform-specific CRISPR models in retinal organoids may provide more mechanistic insights. Integrating isotope tracing with single-cell transcriptomics may also clarify cell type-specific metabolic interactions to delineate the roles of ALT in neurons versus glia.

## Conclusion

This study identifies ALT as an important regulator of metabolic homeostasis in the human neural retina for the first time. We showed that both ALT isoforms are expressed in the retinal tissue, with distinct subcellular localization patterns in Müller glia and photoreceptors, and ALT1 expression is altered in DR. Functional isotope-tracing analyses also demonstrated that retinal ALT is not a passive metabolic marker, but an active node controlling pyruvate-to-alanine flux, nitrogen redistribution, and regional metabolic adaptation across the neural retina.

Importantly, pharmacological ALT inhibition is efficient to suppress alanine synthesis, reshape glutamate-related nitrogen handling, and improve redox balance under oxidative stress. These findings suggest the role of ALT at the intersection of carbon metabolism, amino acid metabolism and antioxidant defense, which processes are centrally involved in retinal vulnerability in diabetes and other metabolic disorders. Although further *in vivo* and isoform-specific studies are desired, our data support that ALT, particularly ALT1-associated metabolic remodeling, may represent a promising therapeutic entry point for diseases characterized by retinal metabolic stress.

## Author Contribution

LZ, MG, JD, TZ, QC and FL conceived the project and designed the experiments. QC, TZ, JZ, MY, SL, FZ, MZ, and MZ performed the experiments, collected data, and conducted the analyses. QC drafted the manuscript, and LZ, MG, JD and FZ critically revised it. All authors read and approved the final version of the manuscript.

## Data availability

Data are available from the authors upon reasonable request.

---

**Figure S1:**
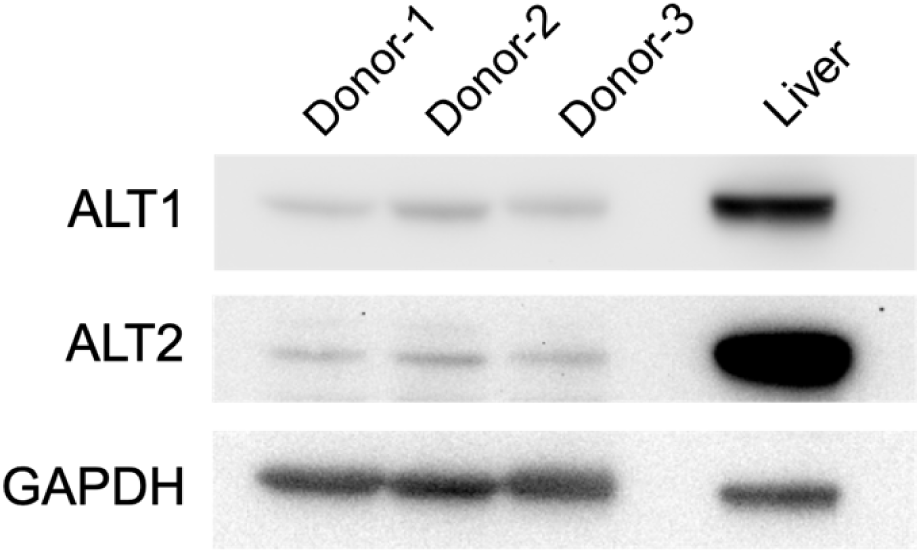
ALT1 and ALT2 are expressed in human retinal explants. Western blot analysis of ALT1 and ALT2 protein expression in human retinal explants. Protein lysates were prepared from 5-mm retinal explants obtained from three individual donors (IDs 0303, 0299, and 0271). Liver tissues were included as positive controls. GAPDH was adopted as the loading control.

**Figure S2.**
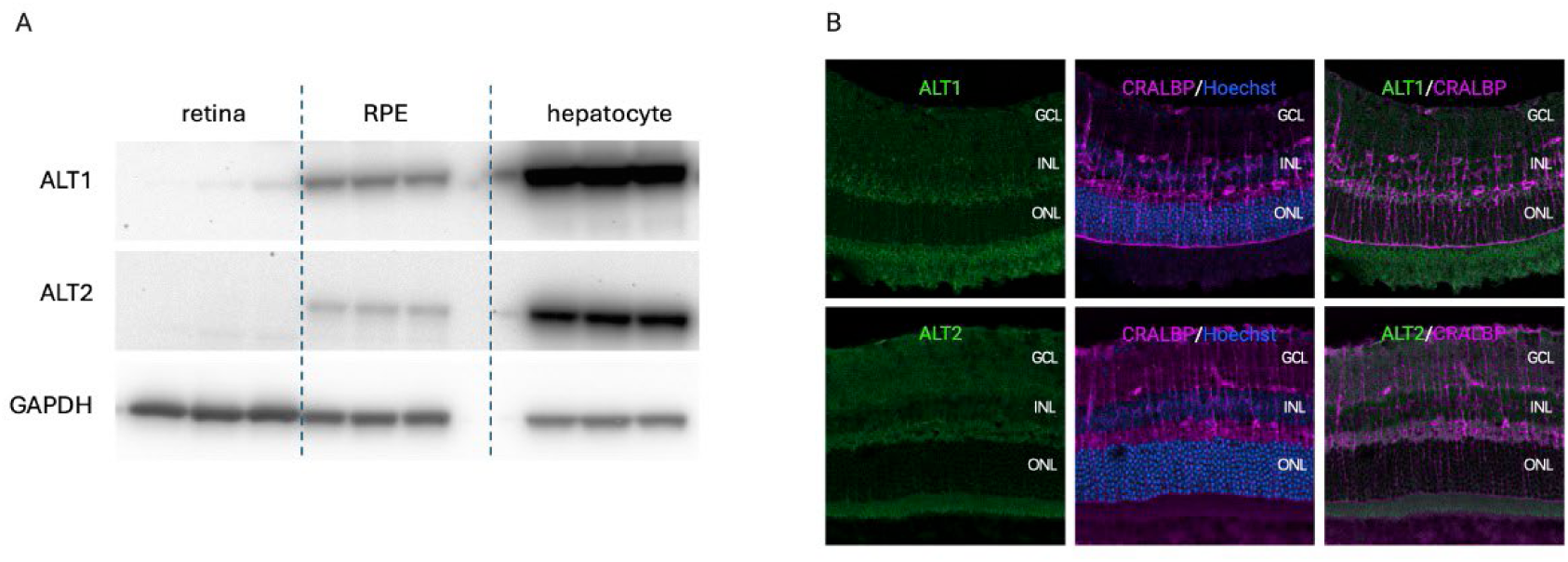
Limited expression and lack of clear cellular localization of ALT1 and ALT2 in mouse retina. A: Western blot analysis of ALT1 and ALT2 protein levels in mouse retina. No distinct or robust ALT1 or ALT2 expression was detected in mouse retinal samples. Hepatocytes were included as positive controls. GAPDH was used as the loading control. B: Immunofluorescence staining of mouse retinal sections for ALT1 or ALT2 (green), co-localization with the Müller glial marker CRALBP (magenta) and nuclear marker Hoechst (blue). ALT1 and ALT2 immunoreactivity was weak and diffuse, without clear enrichment in Müller glia or other defined retinal cell layers.

**Figure S3.**
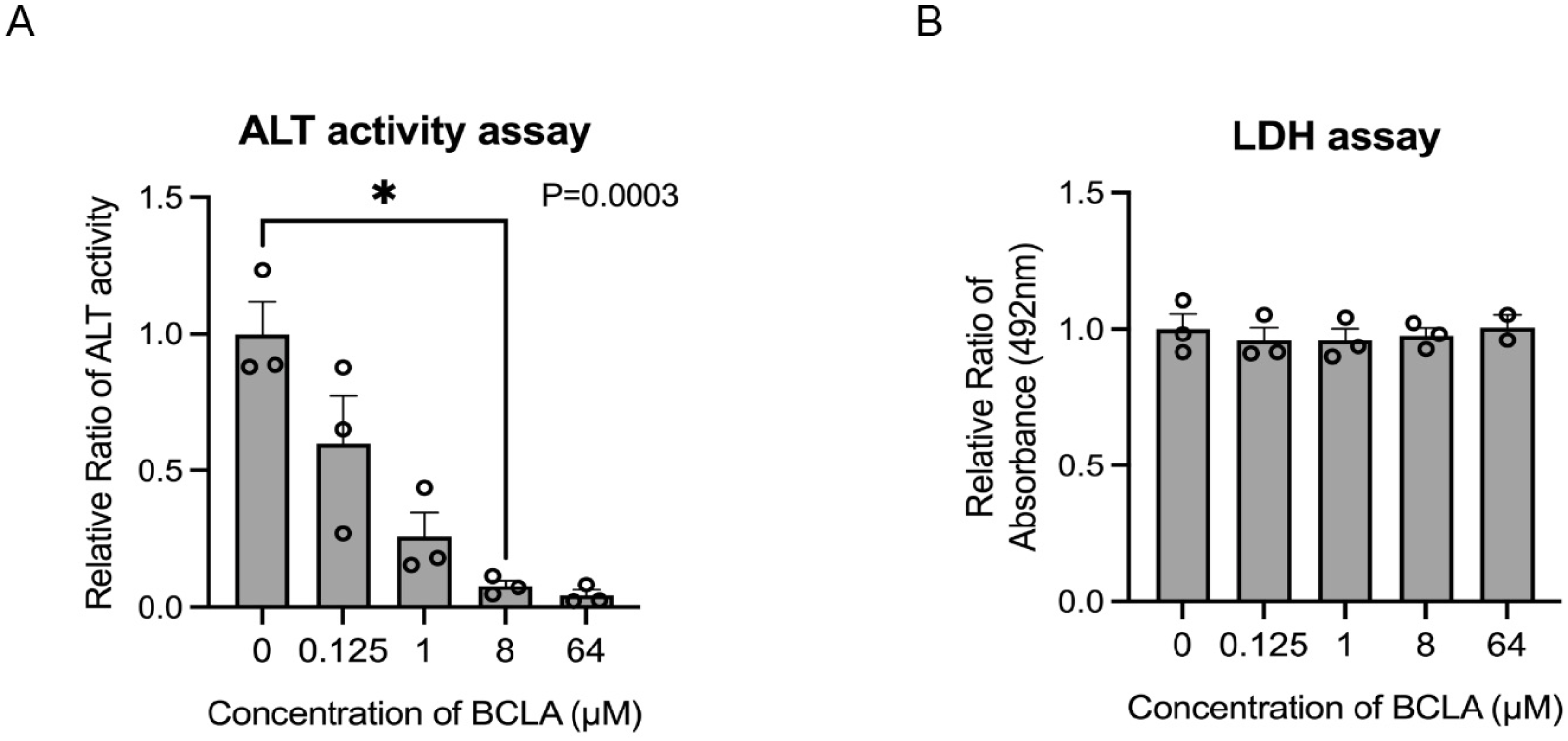
BCLA inhibits ALT activity in human retinal explants without detectable acute cytotoxicity. Human retinal explants from three donors were incubated with increasing concentrations of BCLA (0, 0.125, 1, 8, and 64 μM) for 4 h. A: ALT enzymatic activity was suppressed in a dose-dependent manner following BCLA treatment. B: LDH release showed no significant change across treatment groups, indicating that BCLA did not induce overt acute cytotoxicity under the test conditions. Data are presented as mean ± SE. *, p < 0.05.

**Figure S4:**
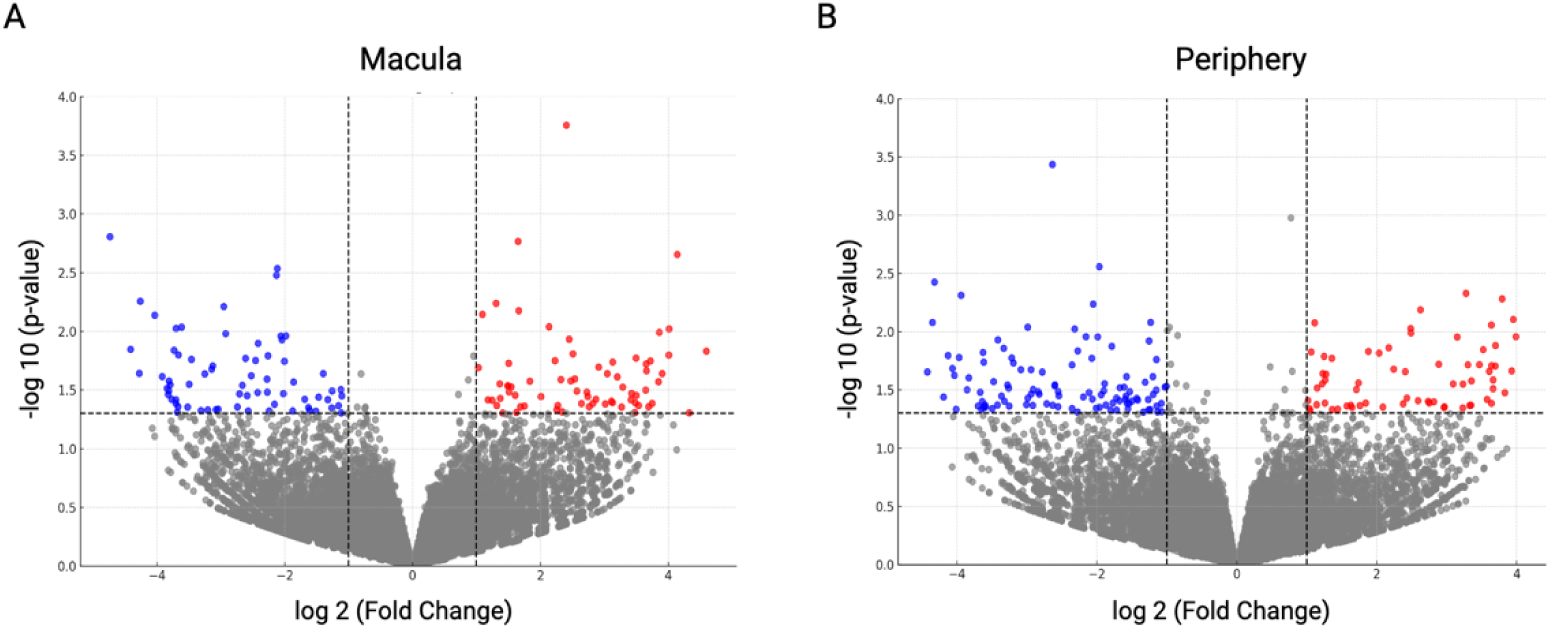
BCLA induces minimal short-term transcriptomic changes in human retinal explants. Macular and peripheral human retinal explants (5 mm diameter) were treated with or without BCLA for 4 h, followed by RNA extraction and bulk RNA sequencing. A& B: Volcano plots showing differentially expressed genes in macular (A) and peripheral (B) retinas. Differential expression was defined using nominal p < 0.05 and |log 2-fold change| > 1. Although a subset of genes showed nominal changes, no genes showed significant changes after multiple testing corrections.

**Figure S5:**
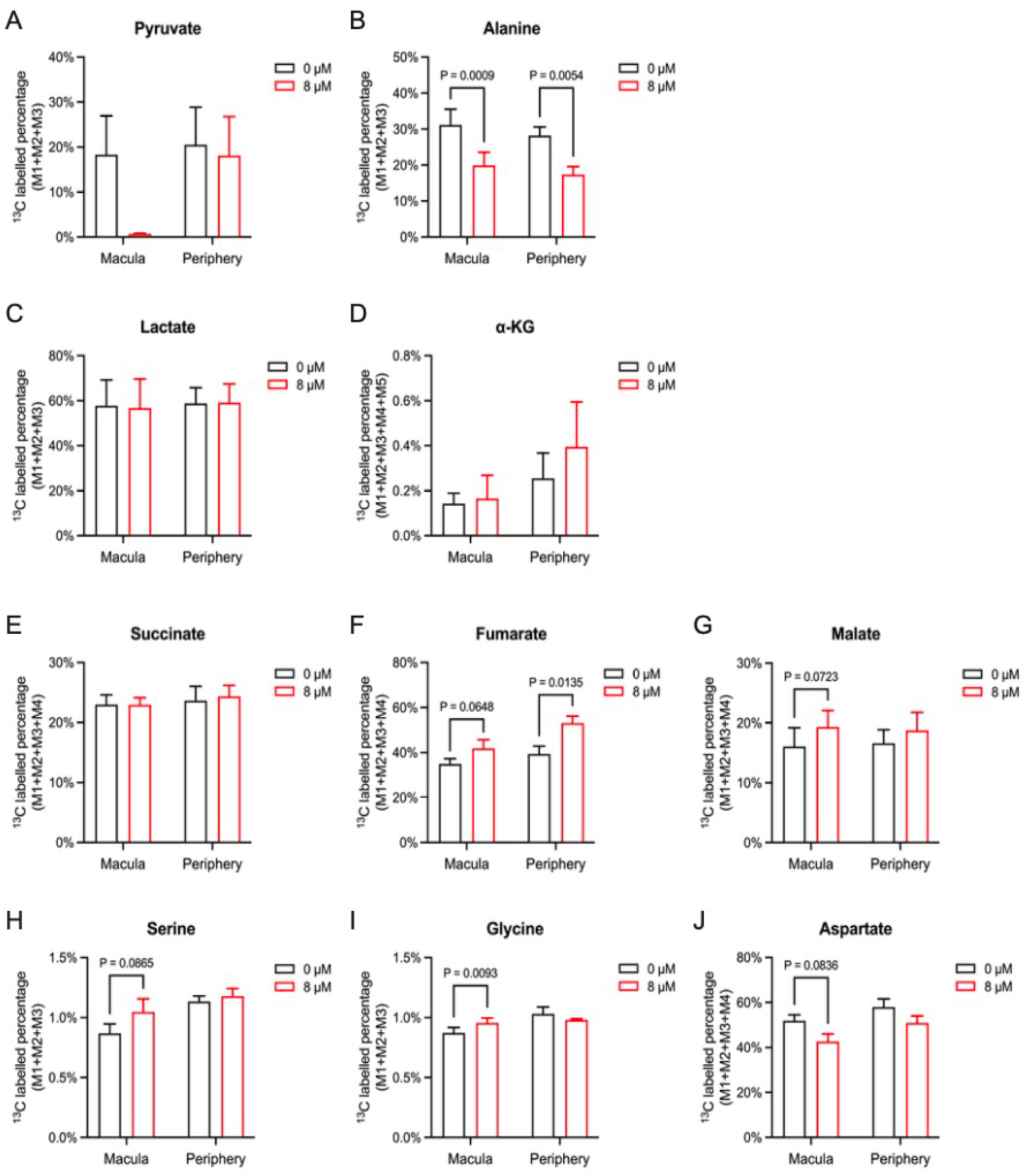
ALT inhibition reduces alanine release and alters extracellular metabolite profiles. A-J: Extracellular metabolite labelling profiles of the control and BCLA-treated explants. BCLA reduced the release of ^13^C-labelled alanine from both macular and peripheral retinas. Additional changes in secreted metabolites suggest region-specific remodeling of extracellular metabolic output following ALT inhibition. Data are presented as mean ± SE.

**Figure S6.**
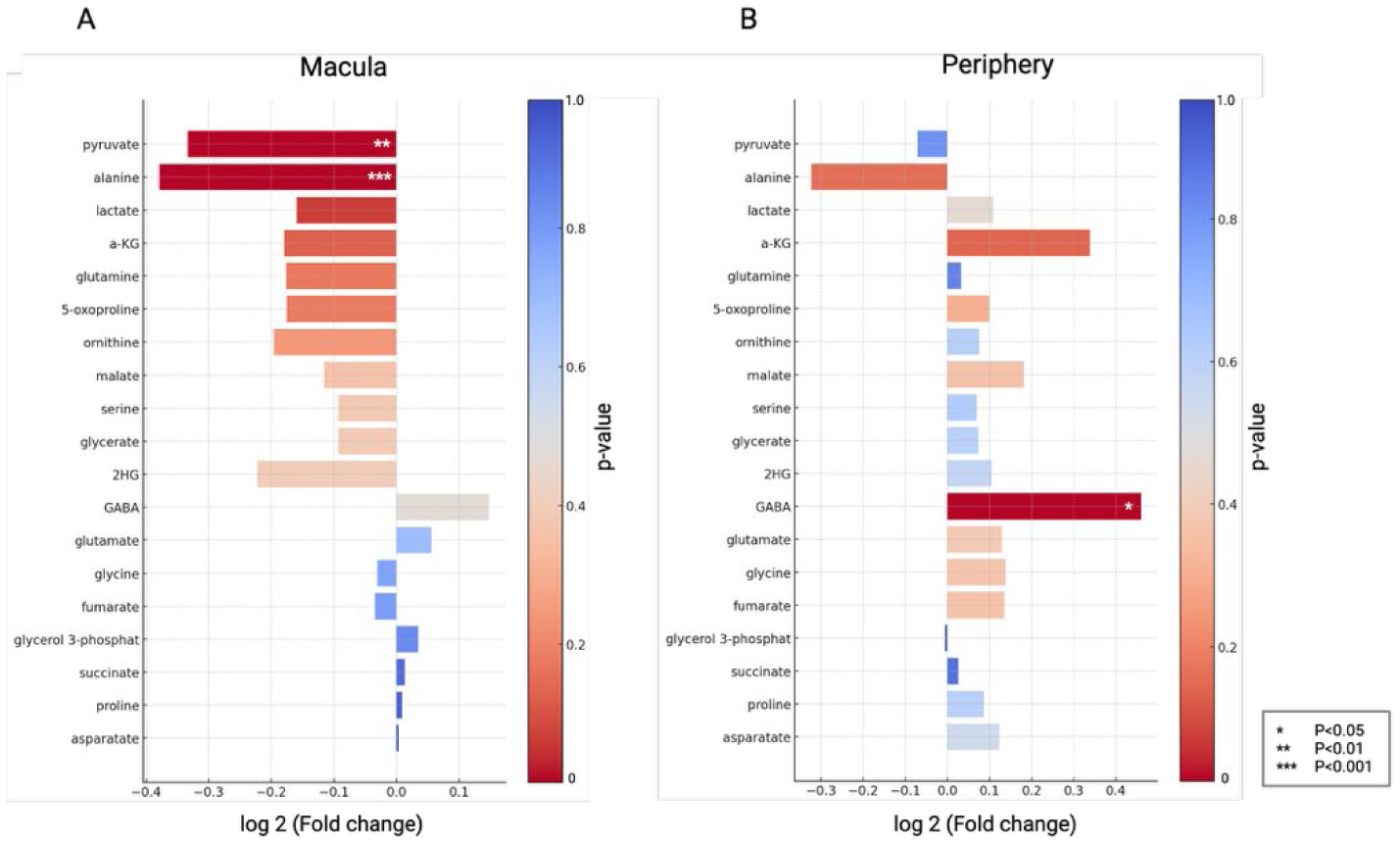
Region-specific metabolic changes in human retinal explants upon BCLA treatment. Bar plots showing log 2-fold changes in total metabolite abundance in macular (left) and peripheral (right) human retinal explants following BCLA treatment. Bars are color-coded as to statistical significance. ALT inhibition produced clear region-specific metabolic responses: alanine and pyruvate were significantly decreased in the macula, whereas several metabolites showed an opposite or compensatory trend in the peripheral retinas. GABA abundance was significantly increased in the periphery retinas.

**Table S1.** Donor information for human retinal explants.

| Donor ID | Gender | Age (year) | Post-mortem Delay (h) | Known eye diseases | Experiments |
| --- | --- | --- | --- | --- | --- |
| D24-1111 | Female | 79 | 28 | No | <sup>13</sup>C-Glucose labelling |
| D24-1115 | Male | 70 | 30 | No |  |
| D24-1142 | Female | 65 | 29 | No |  |
| D24-1189 | Male | 76 | 35 | No |  |
| D24-1304 | Male | 77 | 34 | No |  |
| D24-1192 | Female | 57 | 26 | No | <sup>15</sup>N-Glutamine labelling |
| D24-1218 | Male | 62 | 34 | No |  |
| D24-1238 | Male | 76 | 24 | No |  |
| D24-1277 | Male | 68 | 36 | No |  |
| D24-1306 | Female | 38 | 36 | No |  |
| D24-0863 | Female | 62 | 27 | No | <sup>13</sup>C-Pyruvate labelling |
| D24-0880 | Female | 70 | 24 | No |  |
| D24-0892 | Female | 53 | 36 | No |  |
| D24-0912 | Male | 84 | 18 | No |  |
| D24-0952 | Male | 54 | 34 | No |  |
| D24-0979 | Male | 49 | 36 | No | <sup>13</sup>C-Alanine labelling |
| D24-0988 | Male | 47 | 36 | No |  |
| D24-1039 | Male | 87 | 24 | No |  |
| D24-1066 | Male | 84 | 27 | No |  |
| D24-1149 | Female | 51 | 17 | No |  |

**Table S2.**
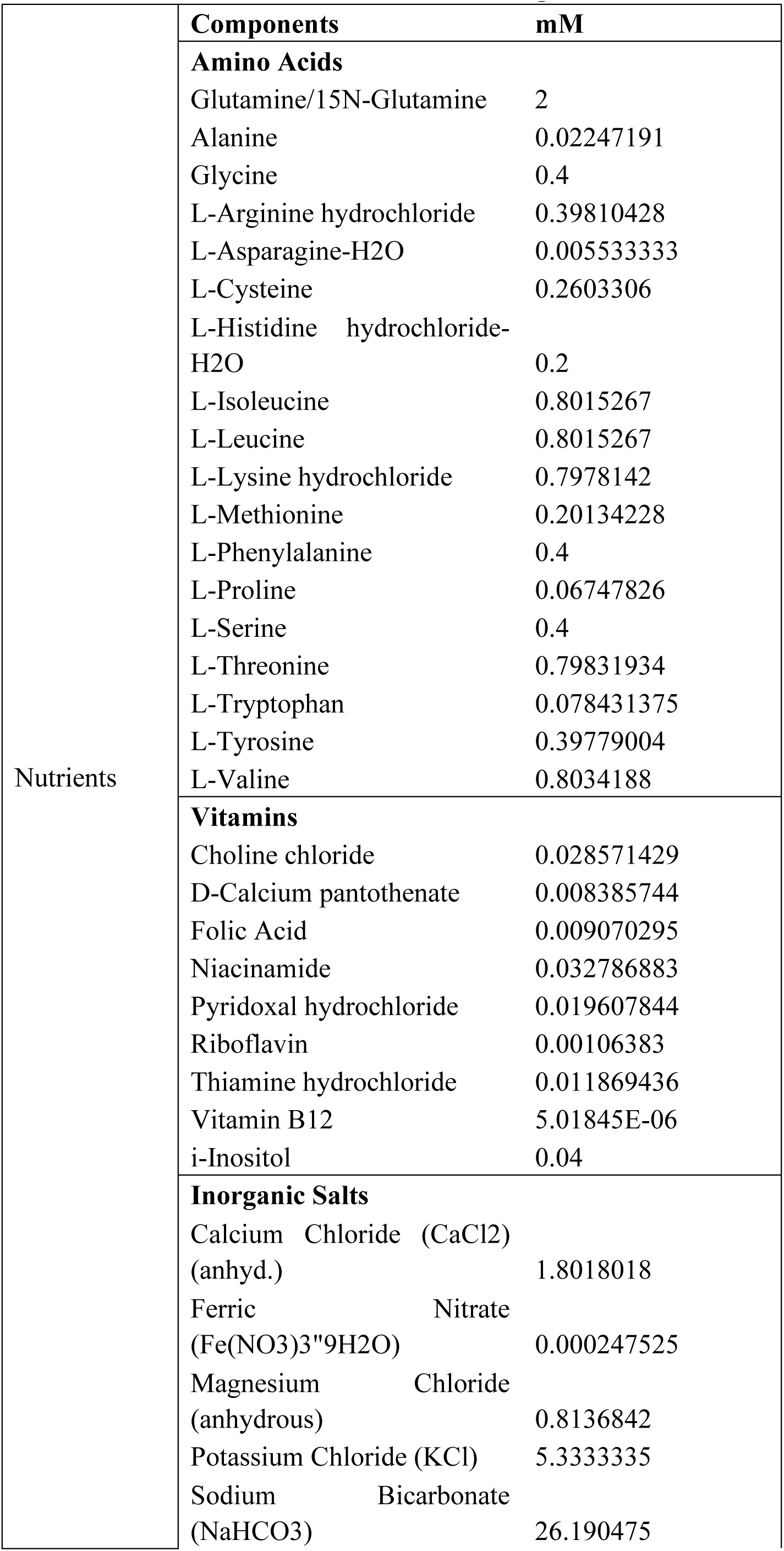
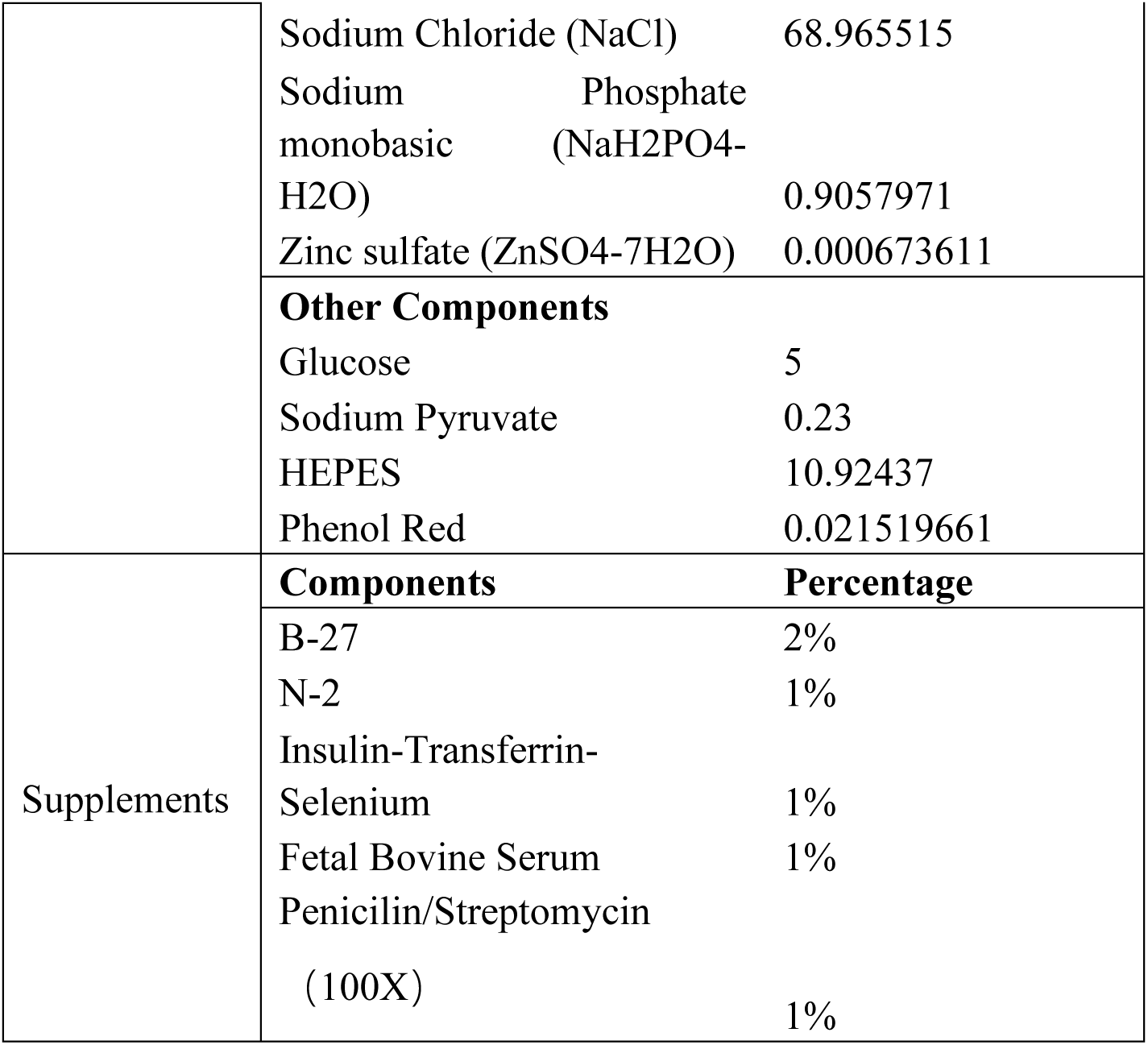
Medium formulation for retinal explants culture.

**Table S3.** The medium of Isotope labelling experiments.

| Group | Medium Composition |
| --- | --- |
| <sup>13</sup>C-glucose | 5mM <sup>13</sup>C-glucose + modified neuronal culture medium without normal glucose |
| <sup>15</sup>N-glutamine | 2mM <sup>15</sup>N-glutamine + modified neuronal culture medium without normal glutamine |
| <sup>13</sup>C-pyruvate | 0.23mM <sup>13</sup>C-pyruvate + modified neuronal culture medium without normal sodium pyruvate |
| <sup>13</sup>C-alanine | 0.02mM <sup>13</sup>C-alanine + modified neuronal culture medium without normal alanine |

